# Machine Learning-Based Bioactivity Prediction of Potential PPAR-*γ* Agonists for the Management of Diabetes

**DOI:** 10.1101/2025.08.09.669467

**Authors:** Juhi Madhwani, Murugesan Sankaranarayanan

## Abstract

This research paper presents a machine learning approach to predict bioactivity of compounds that can act as PPAR-gamma agonist, a critical target for diabetes treatment. Using data from the ChEMBL database, molecular descriptors were calculated and a Random Forest model was developed, achieving an **R**^2^ score of **0.83**. Key molecular features influencing bioactivity were identified, and a web application was created for real-time predictions. This approach demonstrates how computational methods can accelerate drug discovery for diabetes treatment.

## I. Introduction

Peroxisome Proliferator-Activated Receptor Gamma (PPAR *γ*) is a nuclear receptor and transcription factor pivotal in regulating glucose metabolism, lipid storage, and adipocyte differentiation. Primarily expressed in adipose tissue, liver, and skeletal muscle, PPAR *γ* has garnered significant attention in diabetes research due to its critical involvement in insulin sensitivity and glucose homeostasis [1]. With type 2 diabetes mellitus (T2DM) affecting over 460 million people worldwide, the development of effective therapeutic agents targeting PPAR *γ* remains a significant focus in pharmaceutical research.

Traditional drug discovery processes are time-consuming and resource-intensive, often taking 10-15 years and costing billions of dollars to bring a new drug to the market. Computational approaches, particularly machine learning (ML) techniques, offer promising alternatives to accelerate the drug discovery pipeline [2]. By leveraging bioactivity data and molecular descriptors, ML models can predict the activity of novel compounds against specific targets, prioritizing candidates for synthesis and biological testing.

This study aims to develop and validate a machine learning framework for predicting compound bioactivity against PPAR *γ*, potentially accelerating anti-diabetic drug discovery. Specifically, the objectives include:

1. Collection and preprocessing of PPAR *γ* bioactive compounds from the ChEMBL database
2. Calculation of relevant molecular descriptors using appropriate computational tools
3. Development, validation, and evaluation of machine learning models for bioactivity prediction
4. Identification of key molecular features influencing PPAR *γ* binding
5. Deployment of a user-friendly web application for real-time bioactivity predictions

## II. Background and Related Work

### A. PPAR *γ* and Its Role in Diabetes

PPAR *γ* belongs to the PPAR family of nuclear receptors, which includes PPAR *α*, PPAR *δ* (also known as PPAR *β*), and PPAR *γ*. The PPAR *γ* protein structure consists of several domains: an N-terminal A/B domain with AF1 (activation function 1), a C domain containing the DNA-binding domain (DBD), a D domain (hinge region), and an E domain containing the ligand-binding domain (LBD) and AF2 [3].

PPAR *γ* has been targeted pharmacologically using synthetic agonists, most notably the thiazolidinediones (TZDs). These drugs, including pioglitazone and rosiglitazone, activate PPAR *γ* and have demonstrated efficacy in improving glycemic control and insulin sensitivity in T2DM patients. However, TZDs are associated with adverse effects, including weight gain, fluid retention, and increased risk of heart failure, necessitating the discovery of novel PPAR *γ* modulators with improved safety profiles [3].

### B. Computational Approaches in Drug Discovery

Computational methods have revolutionized drug discovery by enabling virtual screening of compound libraries, prediction of pharmacokinetic properties, and structure-based drug design. Machine learning, a subset of artificial intelligence, has emerged as a powerful tool for bioactivity prediction, leveraging patterns in existing data to forecast the activity of novel compounds [4]. Previous studies have applied various machine learning algorithms, including Support Vector Machines (SVMs), Random Forests, and Neural Networks, to predict compound bioactivity against different targets [5]. For PPAR *γ* specifically, machine learning approaches have been utilized to identify potential agonists and partial agonists, although comprehensive frameworks integrating bioactivity prediction with web-based deployment remain limited.

### C. Molecular Descriptors in Drug Design

Molecular descriptors are numerical representations of chemical properties. Key descriptors in drug design include:

1. Topological Polar Surface Area (TPSA): Measures the surface area occupied by polar atoms, influencing membrane permeability and bioavailability.
2. Lipophilicity (LogP): Represents the partition coefficient between octanol and water, indicating a compound’s affinity for lipid environments.
3. Quantitative Estimate of Drug-likeness (QED): Combines multiple molecular properties into a single score assessing overall drug-likeness.
4. Hydrogen Bond Donors and Acceptors: Critical for target binding interactions.
5. Molecular Weight: Influences absorption and distribution properties.

These descriptors, along with structural fingerprints, provide the foundation for machine learning-based bioactivity prediction [6].

## III. Methodology

### A. Data Collection

The study began with identifying PPAR *γ* as the biological target due to its relevance in diabetes treatment. The ChEMBL database, a comprehensive repository of bioactivity data, served as the primary source of information. Data retrieval was performed using the chembl webresource client Python library, enabling direct access to bioactivity data without manual downloads [7].

After identifying the target (CHEMBL235 for human PPAR *γ*), bioactivity data for compounds interacting with PPAR *γ* was retrieved, including essential details such as compound identifiers, molecular structures in SMILES notation, and IC_50_ values representing the compound’s inhibitory activity [7].

**Listing 1:**
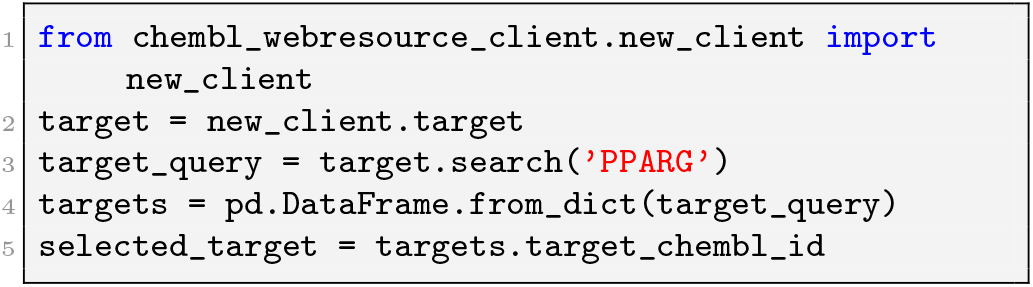
Target identification and data retrieval

### B. Data Pre-processing

The raw data retrieved from ChEMBL required preprocessing to ensure quality and consistency. This involved removing missing values, eliminating duplicates, and standardizing the bioactivity measurements. Compounds were categorized based on their IC_50_ values into three classes [1]:

- Active: IC_50_ *<* 3*µ*M (highly potent compounds)
- Intermediate: IC_50_ 3 to 15*µ*M (moderately active compounds)
- Inactive: IC_50_ *>* 15*µ*M (weakly active or inactive compounds)

Additionally, IC_50_ values were converted to pIC_50_ (−log_10_[IC_50_]) for better numerical handling, as pIC_50_ provides a more normally distributed scale with higher values indicating greater potency [1].

**Listing 2:**
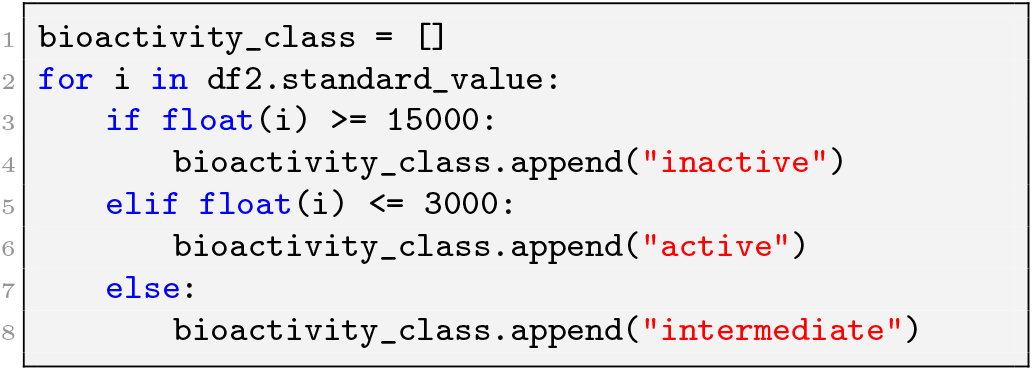
Bioactivity classification

### C. Molecular Descriptor Calculation

Molecular descriptors were calculated to numerically represent the chemical properties of compounds. The RDKit library was used to compute various descriptors including :

1. Topological Polar Surface Area (TPSA)
2. Lipophilicity (ALogP)
3. Number of Hydrogen Bond Donors (HBD)
4. Number of Hydrogen Bond Acceptors (HBA)
5. Fraction of sp3 Hybridized Carbons (FSP3)
6. Quantitative Estimate of Drug-likeness (QED)
7. Number of Rotatable Bonds (NRot)

Additionally, PubChem fingerprints were generated to capture structural features of the molecules. The following code snippet demonstrates the calculation of key descriptors [8]:

**Listing 3:**
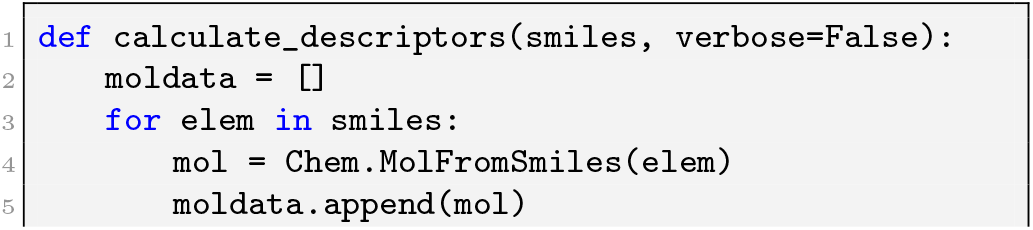
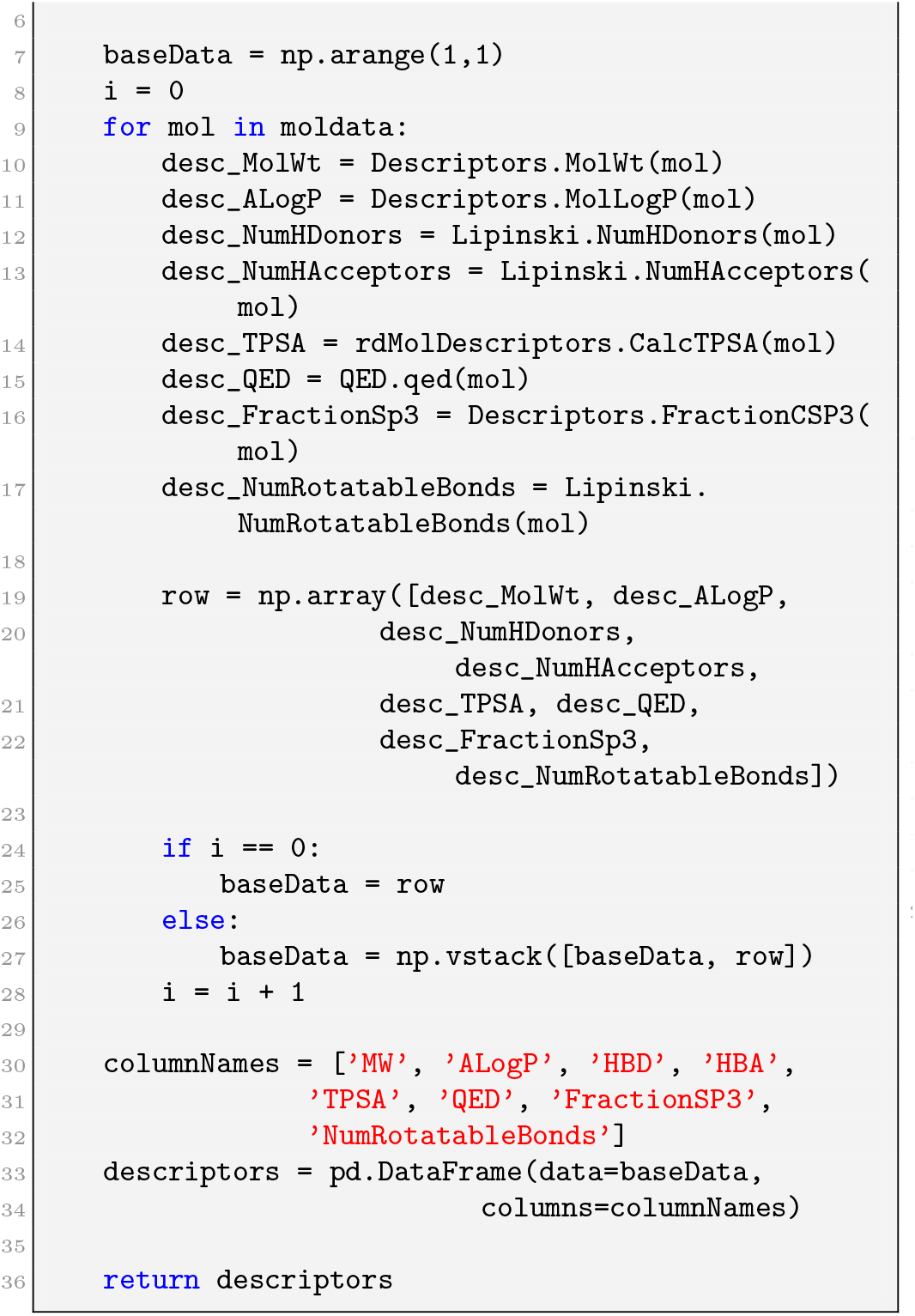
Molecular descriptor calculation

### D. Exploratory Data Analysis (EDA)

Exploratory Data Analysis (EDA) was performed to understand the relationships between molecular descriptors and bioactivity classes. Visualizations, including boxplots and scatterplots, were created to observe how descriptors like molecular weight, TPSA, and LogP vary across active, intermediate, and inactive compounds [1].

Statistical tests, particularly Mann-Whitney U tests, were applied to assess whether the descriptor distributions differed significantly between active and inactive classes, providing insights into which properties are most relevant to bioactivity.

**Listing 4:**
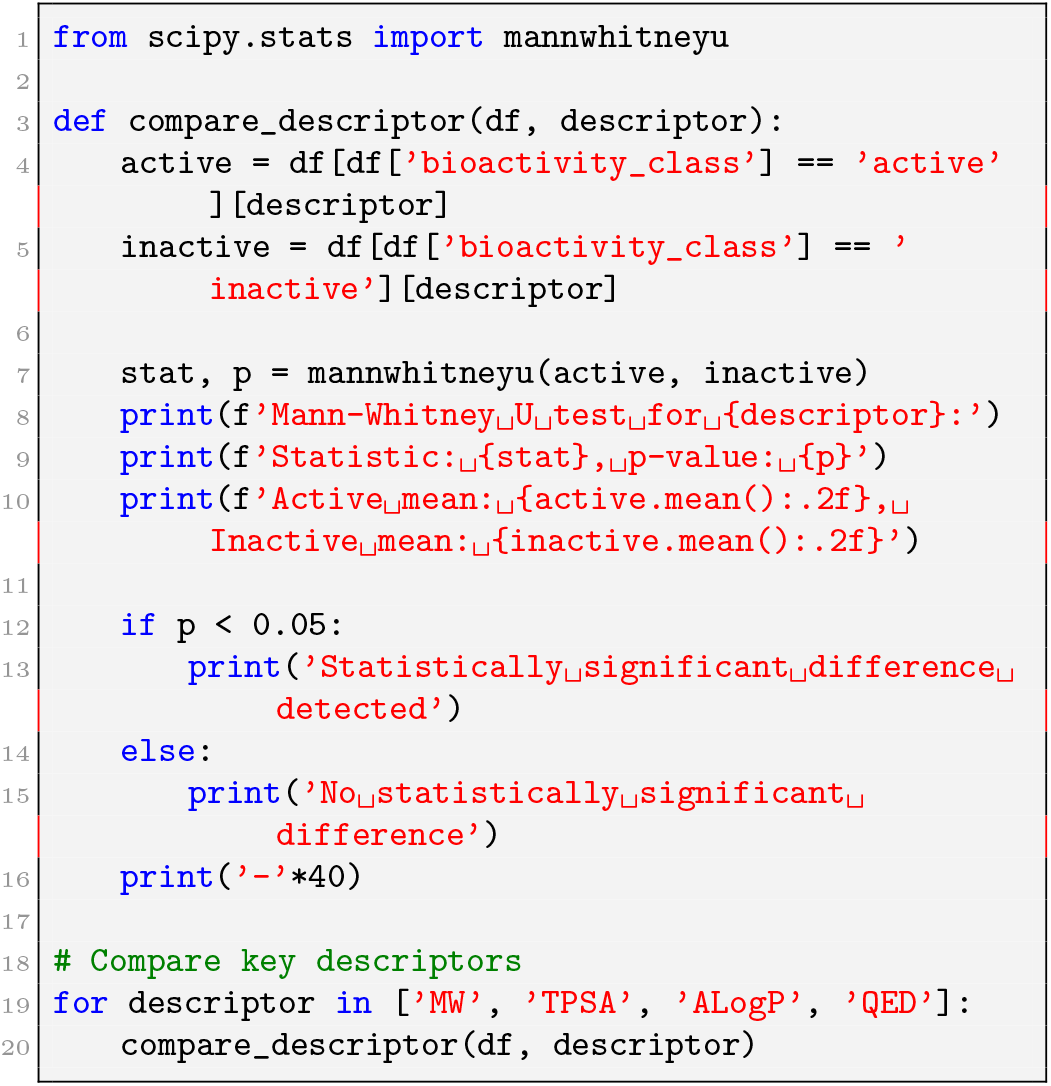
Statistical comparison of descriptors

### E. Model Development

A Random Forest regressor was chosen for predicting pIC_50_ values due to its ability to handle complex, non-linear relationships between descriptors and bioactivity. The model development process included [7,8]:

1. Splitting the dataset into training (80%) and test (20%) sets:
2. Training the model on the training set:
3. Evaluating the model on the test set using metrics such as *R*^2^ and RMSE:
4. Cross-validation to ensure model robustness:
5. Feature importance analysis to identify key molecular descriptors:

**Listing 5:**
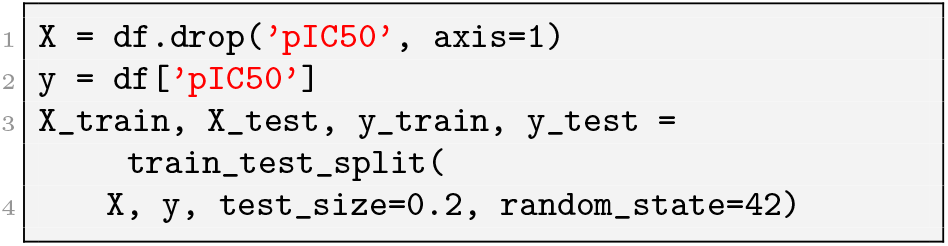
Data splitting

**Listing 6:**
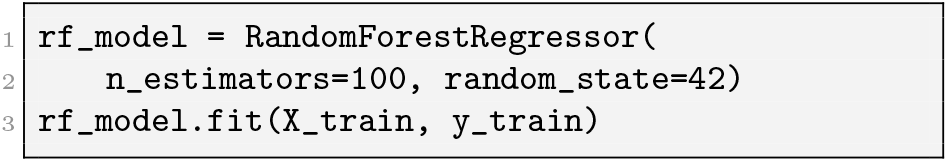
Model training

**Listing 7:**
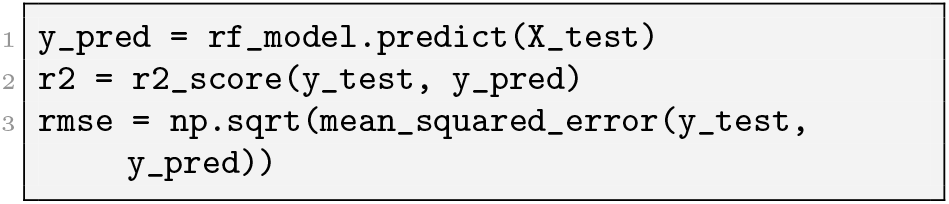
Model evaluation

**Listing 8:**
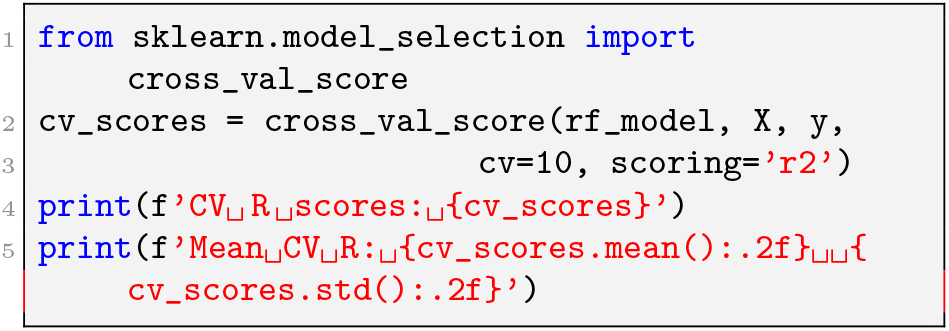
Cross-validation

**Listing 9:**
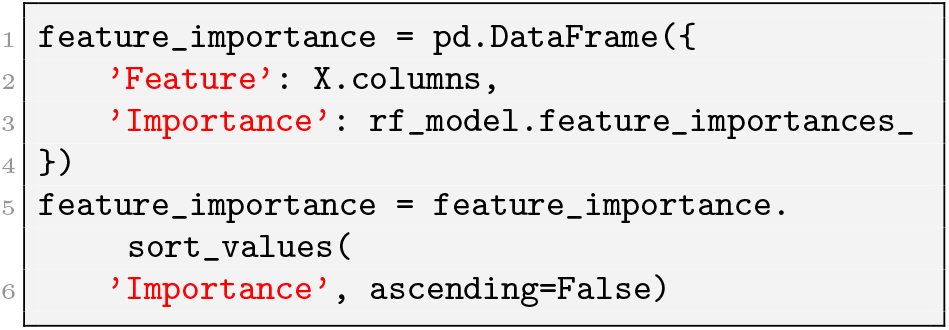
Feature importance analysis

### F. Web Application Development

The trained model was deployed as a web application using Streamlit, a Python library for creating interactive web applications. The application enables users to input molecular structures (as SMILES notation) and receive real-time predictions of bioactivity against PPAR *γ* [7,8].

Key features of the web application include:

1. Input field for SMILES notation
2. Calculation of molecular descriptors for the input compound
3. Prediction of pIC_50_ values using the trained model
4. Classification of compounds as active, intermediate, or inactive

The following code snippet demonstrates the core functionality:

**Listing 10:**
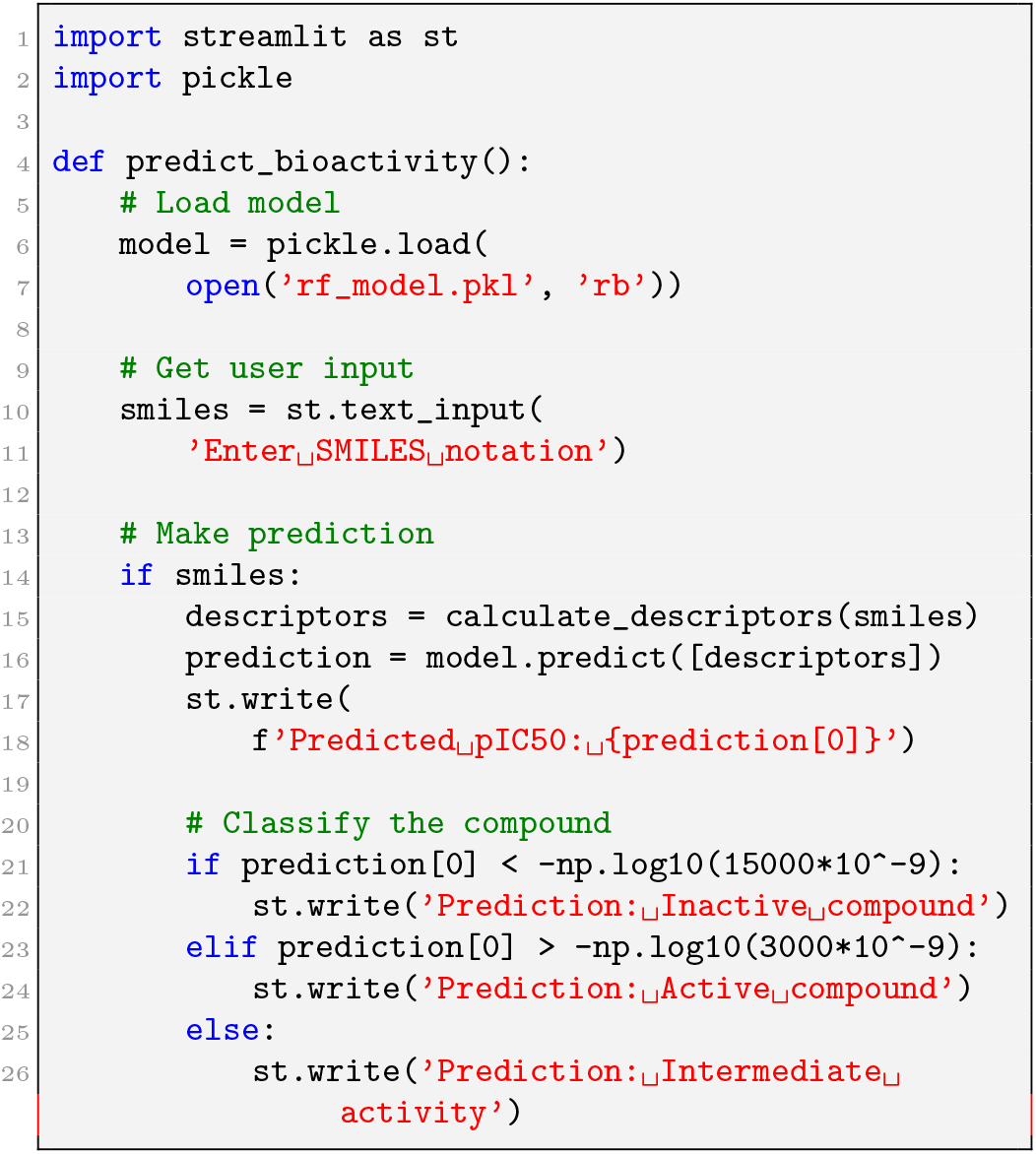
Web application implementation

## IV. Results

### A. Exploratory Data Analysis (EDA) Findings

The exploratory data analysis revealed significant differences in molecular descriptors between active and inactive compounds.[1]:

Topological Polar Surface Area (TPSA): Active compounds generally showed lower TPSA values (mean 75 Å^2^) compared to inactive compounds (mean 100 Å^2^), suggesting that lower polar surface area enhances permeability across cell membranes and potentially improves binding to :

Topological Polar Surface Area (TPSA): Active compounds generally showed lower TPSA values (mean 75 Å^2^) compared to inactive compounds (mean 100 Å^2^), suggesting that lower polar surface area enhances permeability across cell membranes and potentially improves binding.

Lipophilicity (ALogP): Active compounds exhibited moderate ALogP values (range: 2.5-7.5), indicating a balance between hydrophobicity and hydrophilicity. Inactive compounds often had values outside this optimal range, being either too hydrophilic or too hydrophobic.

Molecular Weight (Mol wt): Active compounds showed a narrower distribution of molecular weights compared to inactive compounds, with most active compounds falling within the range of 350-550 Da.

Quantitative Estimate of Drug-likeness (QED): Active compounds generally displayed higher QED values, suggesting better overall drug-like properties..

Lipophilicity (ALogP): Active compounds exhibited moderate ALogP values (range: 2.5-7.5), indicating a balance between hydrophobicity and hydrophilicity. Inactive compounds often had values outside this optimal range, being either too hydrophilic or too hydrophobic.

Molecular Weight (Mol.wt): Active compounds showed a narrower distribution of molecular weights compared to inactive compounds, with most active compounds falling within the range of 350-550 Da.

Quantitative Estimate of Drug-likeness (QED): Active compounds generally displayed higher QED values, suggesting better overall drug-like properties.

Fig. 1 shows the distribution of compounds across bioactivity classes, with active compounds comprising a substantial portion of the dataset, which ensures balanced model training.

**Fig. 1.**
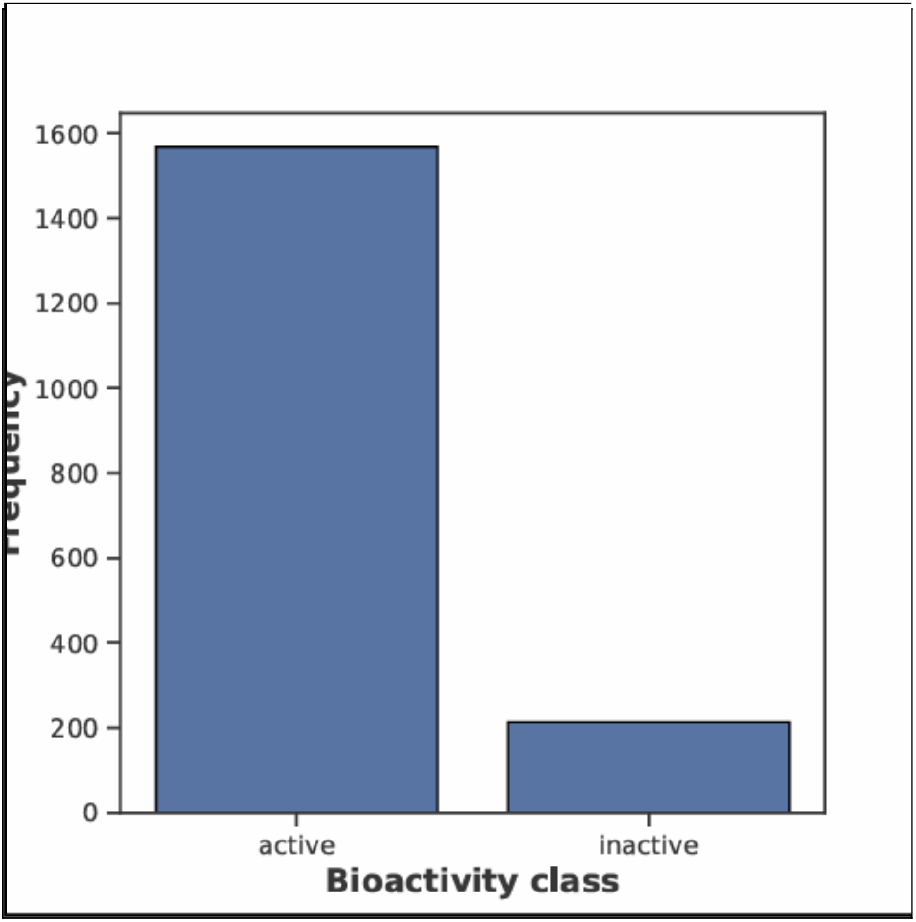
Distribution of compounds across bioactivity classes (Active: IC_50_ ¡3M, Intermediate: 3-15M, Inactive: ¿15M). The balanced distribution supports robust model training by preventing class imbalance issues.

Fig. 2 illustrates the distribution of pIC_50_ values across different bioactivity classes, showing a clear separation between active and inactive compounds:

**Fig. 2.**
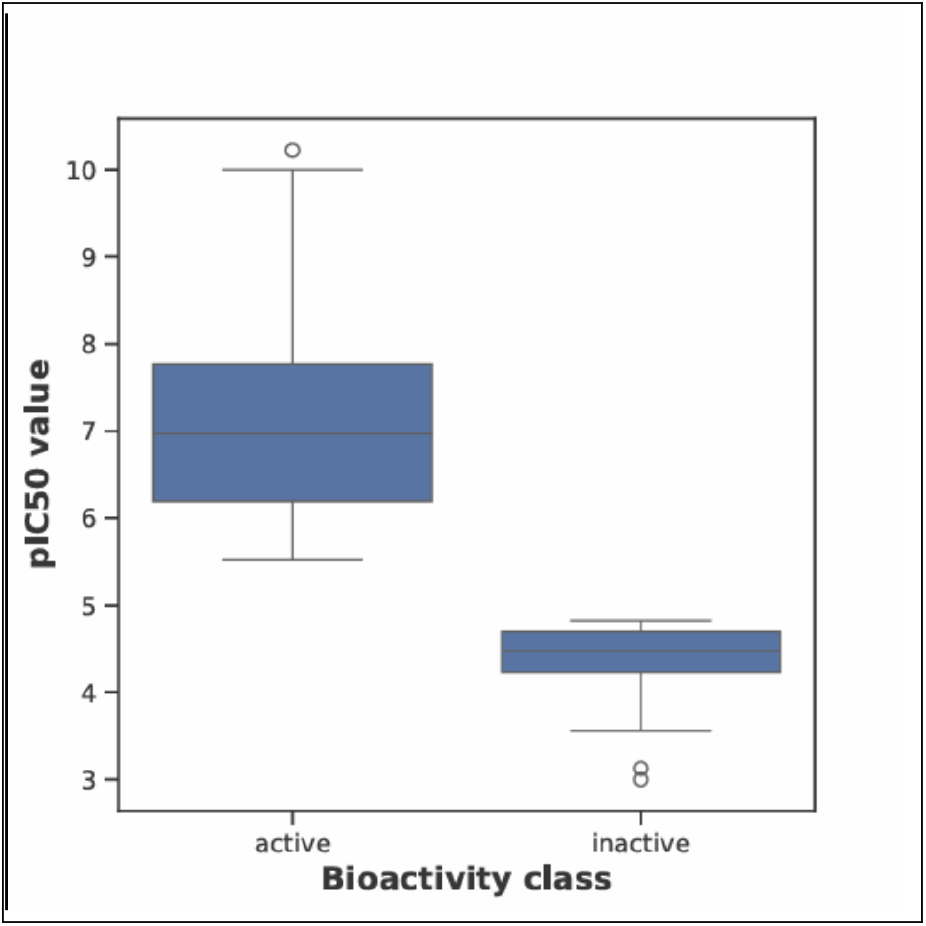
Distribution of pIC_50_ values. Active compounds (bioactivity class) show significantly higher pIC_50_ values, indicating greater potency against PPAR-*γ*. The distinct separation between active and inactive compounds supports reliable classification.

Analysis of specific molecular descriptors revealed important structure-activity relationships (Fig. 3 and Fig. 4):

**Fig. 3.**
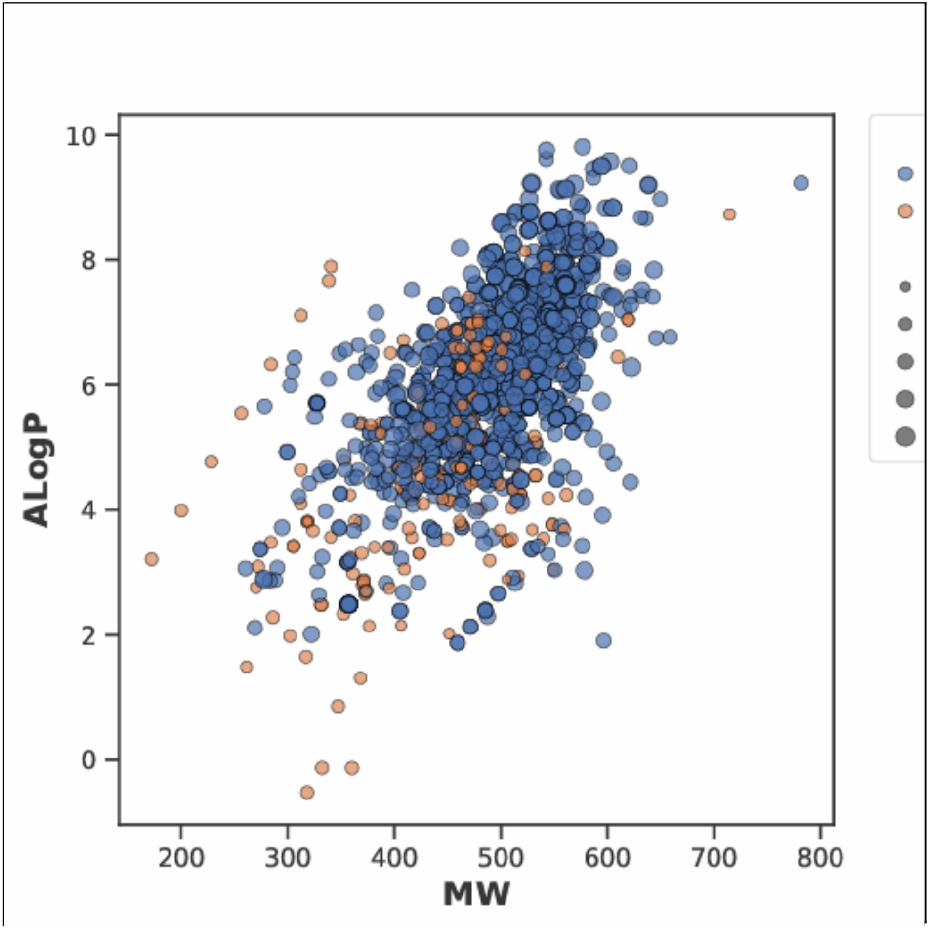
Relationship between Molecular Weight (MW) and Lipophilicity (ALogP) colored by bioactivity class. Active compounds (blue points) cluster primarily in the MW range of 350-550 Da and ALogP range of 3.5-5.5, suggesting optimal physicochemical properties for PPAR-*γ* binding.

**Fig. 4.**
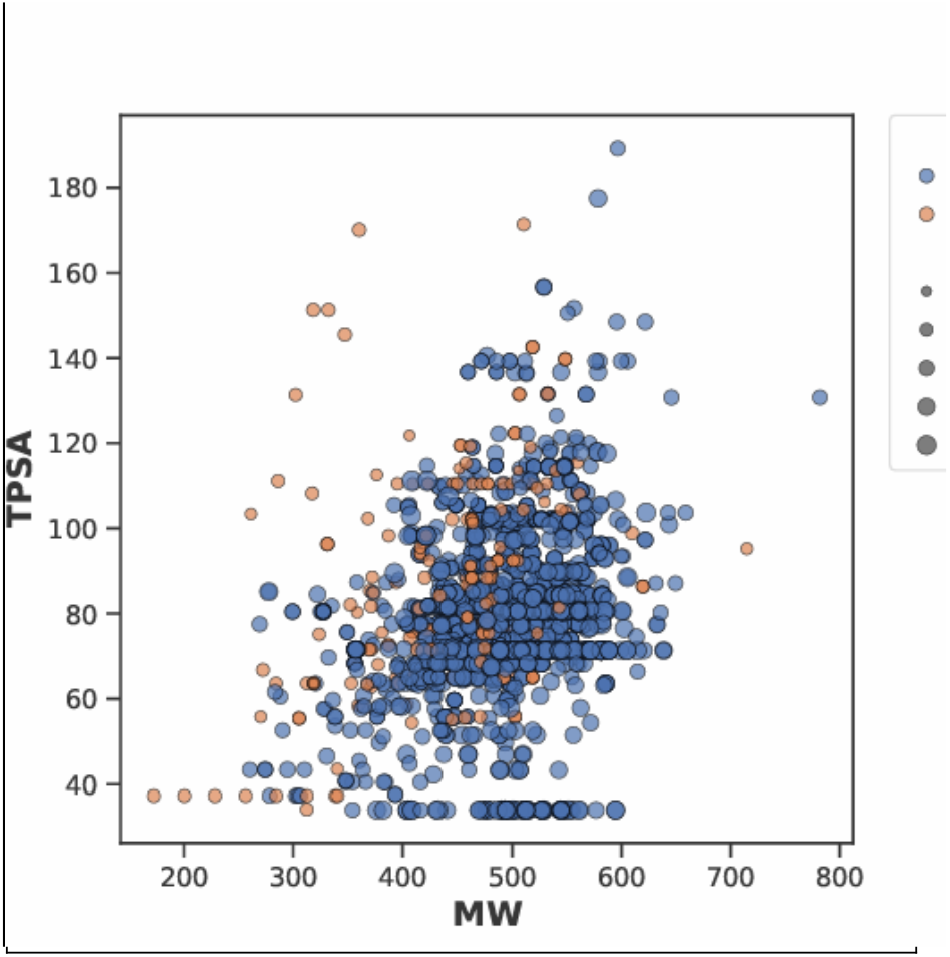
Correlation between Molecular Weight (MW) and Topological Polar Surface Area (TPSA). Active compounds (blue) predominantly exhibit TPSA values below 100Å^2^, regardless of molecular weight, indicating the importance of limited polarity for effective PPAR-*γ* activity.

Statistical analysis using Mann-Whitney U tests confirmed that these differences were statistically significant (*p <* 0.05), supporting the relevance of these descriptors in discriminating between active and inactive compounds [7].

### B. Model Performance

The Random Forest regressor demonstrated strong predictive performance :

- R^2^ Score: 0.83, indicating that 83% of the variance in pIC_50_ values could be explained by the model.
- Root Mean Squared Error (RMSE): 0.27, showing relatively small prediction errors.
- Mean Absolute Error (MAE): 0.21, further confirming the model’s accuracy.

Fig. 5 illustrates the correlation between predicted and experimental pIC_50_ values, with points closely aligned along the identity line (red), confirming the model’s predictive accuracy across the full bioactivity range (pIC_50_ 3-10).

**Fig. 5.**
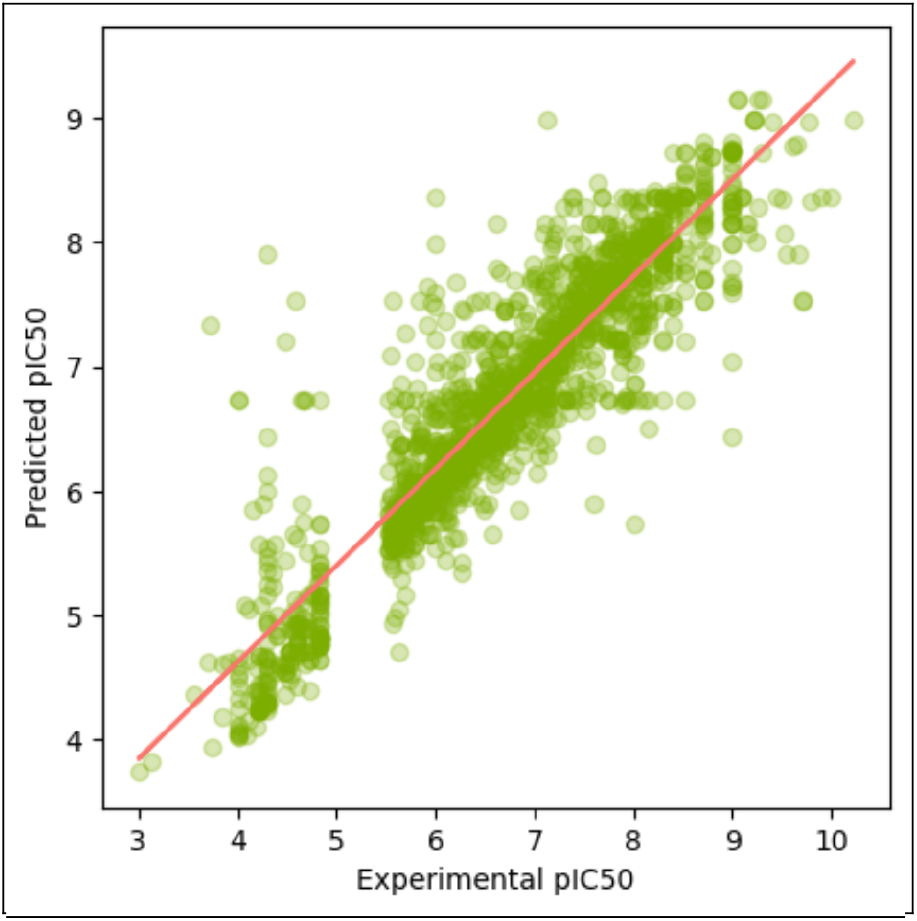
Scatter plot showing correlation between experimental and predicted pIC_50_ values. The strong alignment along the diagonal (red line) demonstrates high predictive accuracy (R^2^ = 0.83). Note the consistent performance across both low (3-5) and high (7-9) bioactivity ranges, with minimal systematic bias.

Feature importance analysis revealed that Pub-Chem fingerprints, TPSA, ALogP, and QED were among the most influential predictors of PPAR *γ* bioactivity, aligning with the findings from the exploratory data analysis.

### C. Web Application Functionality

The Streamlit web application successfully provided real-time bioactivity predictions [7,8]:

1. Users could input SMILES notation for compounds of interest.
2. The application calculated molecular descriptors and generated PubChem fingerprints on-the-fly.
3. The trained Random Forest model predicted pIC_50_ values and classified compounds as active, intermediate, or inactive.
4. Results were displayed in an intuitive format, including the predicted pIC_50_ value and the bioactivity classification.
5. Visualizations compared the input compound’s descriptors to the distribution in the training dataset.

The application demonstrated the practical applicability of the developed model for virtual screening and prioritization of compounds for synthesis and testing [7,8].

## V. Discussion

### A. Interpretation of Findings

The results highlight several key insights regarding molecular features that influence PPAR *γ* binding:

1. Lower TPSA values in active compounds (Fig. 4) suggest that reduced polarity enhances membrane permeability and potentially facilitates interaction with the predominantly hydrophobic binding pocket of PPAR *γ* [1].
2. The optimal range of lipophilicity (ALogP) for active compounds (Fig. 3) aligns with previous studies on PPAR *γ* agonists, which typically feature both hydrophobic regions for interaction with the ligand-binding domain and hydrophilic groups for hydrogen bonding.
3. The importance of PubChem fingerprints in the predictive model indicates that specific structural features, beyond simple physicochemical properties, play crucial roles in determining bioactivity.
4. The strong predictive performance of the Random Forest model (Fig. 5) suggests that the complex, non-linear relationships between molecular descriptors and PPAR *γ* binding are effectively captured by this ensemble learning approach.

### B. Comparison with Previous Studies

The findings align with previous computational studies on PPAR *γ* ligands, which have similarly identified lipophilicity and hydrogen-bonding capabilities as critical determinants of binding affinity. However, this study extends beyond earlier work by:

1. Incorporating a comprehensive set of molecular descriptors and structural fingerprints.
2. Developing a high-performing machine learning model specifically trained on PPAR *γ* bioactivity data.
3. Deploying the model as an accessible web application for real-time predictions.
4. Providing a workflow that can be adapted to other therapeutic targets.

Compared to traditional docking-based virtual screening, this machine learning approach offers advantages in terms of computational efficiency and the ability to capture nonlinear structure-activity relationships [5].

### C. Limitations and Future Directions

Despite the promising results, several limitations should be acknowledged:

1. The model was trained on in vitro bioactivity data, which may not fully reflect in vivo efficacy and safety profiles.
2. The dataset size, while substantial, could be expanded to enhance predictive performance further.
3. The current model focuses solely on PPAR *γ* and does not account for potential off-target interactions or ADMET properties.

Future work could address these limitations through:

1. Integration of additional endpoints, including ADMET properties and off-target effects.
2. Exploration of deep learning approaches, particularly graph neural networks, which can directly operate on molecular graphs.
3. Incorporation of structural information about PPAR *γ* through hybrid machine learning-molecular docking approaches.
4. Experimental validation of predicted high-activity compounds to verify the model’s practical utility.

## VI. Conclusion

This study successfully developed and deployed a machine learning framework for predicting compound bioactivity against PPAR *γ*, a key target for anti-diabetic drugs. The Random Forest model achieved strong predictive performance (R^2^ = 0.83, RMSE = 0.27), demonstrating its potential utility in virtual screening and lead optimization.

Key molecular descriptors influencing PPAR *γ* binding were identified, including TPSA, ALogP, and specific structural fingerprints, providing insights for rational design of novel PPAR *γ* modulators. The web application deployment demonstrates the practical applicability of the approach, enabling researchers to make real-time predictions without specialized computational expertise.

This framework can accelerate the discovery of novel antidiabetic compounds by prioritizing candidates for synthesis and biological testing, potentially reducing the time and resources required for drug discovery and development. Additionally, the methodology is generalizable and can be adapted to other therapeutic targets beyond PPAR *γ*.

Future work will focus on expanding the model’s capabilities, incorporating additional endpoints, exploring advanced machine learning architectures, and experimental validation of predicted high-activity compounds.

## ACKNOWLEDGMENT

The authors thank the Birla Institute of Technology and Science, Pilani, Pilani Campus for providing the necessary infrastructure and computational resources. This work utilizes methodologies and code frameworks developed by Professor Chanin Nantasenmat (Data Professor) through his educational content, which has been instrumental in implementing the machine learning and web application components of this study.

